# A Rho-GTPase based model explains spontaneous collective migration of neural crest cell clusters

**DOI:** 10.1101/181743

**Authors:** Brian Merchant, Leah Edelstein-Keshet, James J. Feng

## Abstract

We propose a model to explain the spontaneous collective migration of neural crest cells in the absence of an external gradient of chemoattractants. The model is based on the dynamical interaction between Rac1 and RhoA that is known to regulate the polarization, contact inhibition and co-attraction of neural crest cells. Coupling the reaction-diffusion equations for active and inactive Rac1 and RhoA on the cell membrane with a mechanical model for the overdamped motion of membrane vertices, we show that co-attraction and contact inhibition cooperate to produce persistence of polarity in a cluster of neural crest cells by suppressing the random onset of Rac1 hotspots that may mature into new protrusion fronts. This produces persistent directional migration of cell clusters in corridors. Our model confirms a prior hypothesis that co-attraction and contact inhibition are key to spontaneous collective migration, and provides an explanation of their cooperative working mechanism in terms of Rho GTPase signaling. The model shows that the spontaneous migration is more robust for larger clusters, and is most efficient in a corridor of optimal confinement.

## 1 Introduction

During vertebrate embryogenesis, neural crest cells (NCCs) delaminate from the neural plate, become highly migratory through an epithelial-to-mesenchymal transition, and then travel long distances to target locations where they differentiate into a wide range of cell types. To coordinate this long range migration, NCCs must integrate information from a variety of external signals, including chemoattractants [1–4]. The process is highly complex, with apparently different mechanisms among different species and between cranial and trunk streams of NCC [5], and thus many unanswered questions remain. Surprisingly, a body of evidence suggests that NCCs may migrate *spontaneously* as a group in the absence of chemoattractants. *In vitro* experiments, with NCCs plated at the end of a fibronectin corridor, find that the NCCs are able to migrate down the corridor with high persistence, in the absence of any directional information from an external chemoattractant [6]. Spontaneous collective motion has also been documented for clusters of bovine capillary endothelial cells [7] and epithelial sheets of Madin-Darby canine kidney (MDCK) cells [8] in confined geometries.

*In vivo*, before the budding of endodermal pouches in zebrafish, NCCs require the presence of chemoattractant Sdf1, sensed through filopodia, to direct their collective migration. However, after the budding of endodermal pouches, NCCs are able to migrate efficiently even when their filopodia are strongly antagonized by the F-actin depolymerizing drug Latrunculin B [9]. The pouches apparently provide sufficient physical confinement as guidance. Other *in vivo* experiments have shown that groups of enteric NCCs transplanted to their target location are able to migrate in the reverse direction, suggesting the absence of an external guiding chemoattractant gradient [10]. Instead, a spontaneous symmetry-breaking appears to determine the direction of migration.

The current hypothesis for this spontaneous collective directional migration is that it emerges from two intercellular interactions between NCCs: contact inhibition of locomotion (CIL) and co-attraction (COA). CIL describes the tendency of NCCs to move away from each other upon contact, a process mediated by N-cadherins and the non-canonical Wnt signalling pathway [11, 12]. COA describes how NCCs attract each other through the autocrine production of the short-ranged chemoattractant C3a and its receptor C3aR [6]. Two recent computational models [13, 14] have sought to demonstrate how spontaneous migration of a group of NCCs can arise from the simultaneous action of CIL and COA. The model of Woods *et al.* [13] treats the cells as particles that move according to Newton’s second law of motion and interact according to rules that represent COA and CIL. Later, the cellular Potts model of Szabo *et al.* [14] avoids the inertia-based second-order dynamics of [13], and reproduces CIL and COA through lattice-based rules that govern the preferential direction of motion for the entire cell. While these models successfully reproduce the spontaneous persistent migration of cell clusters, they leave a fundamental question unanswered: how do COA and CIL, which have no inherent anisotropy, cause a cluster of cells to develop a collective polarity and persistent movement in a certain direction? More specifically for the essentially one-dimensional (1D) corridors tested in these models, how do rules of COA and CIL produce a symmetry-breaking between left- and rightward migration? These questions have motivated the present study.

We started out by building a simple model that produced CIL and COA (more details in the Supporting Information). For the current purpose, suffices it to say that in this model, CIL and COA alone failed to reproduce spontaneous migration that persisted in direction. In the corridor geometry of Szabo *et al.* [14], a cluster of model cells exhibited CIL and COA and remained cohesive. But its centroid merely executed a random walk to the left and the right. There was no symmetry-breaking and no persistent directional migration.

This led us to reexamine the earlier models for additional mechanisms that may have helped CIL and COA to produce persistent directionality in collective migration. In [13], one such candidate mechanism is the inertia of the particles. In reality, cell migration is non-inertial and dominated by overdamped dynamics. The cellular Potts model [14, 15] requires the polarization of a cell to be biased in favor of the most recent displacement. An additional rule explicitly enhances persistence of polarization in the presence of neighbors “to achieve realistic persistence of free cells and cells in clusters”. From the above, we hypothesize that aside from CIL and COA, these models required a third ingredient—persistence of polarity (POP) from one time step to the next—in order to reproduce spontaneous directional cell migration in clusters. This seems similar to numerous active-particle models that rely on particle-level rules of alignment to produce symmetry-breaking and pattern formation in collective motion [16, 17].

Short-term persistence of directional motion occurs naturally as, for one, the remodeling of the cytoskeleton takes tens of seconds. Nevertheless, invoking postulated rules of POP seems unsatis-factory to us as an explanation for spontaneous collective migration of NCCs over the timescale of hours. This issue partly motivated our work. Instead of a rule-based implementation of CIL and COA as found in earlier models [13, 14], we wished to base both on known biochemical pathways of key regulators such as Rac1 and RhoA [6, 18]. Furthermore, we asked whether and how POP could result from such biochemical pathways, and, if so, how it would interact with CIL and COA chemically or mechanically to produce spontaneous collective migration of NCCs.

In this paper, we present a multi-cell two-dimensional (2D) model for spontaneous directional migration of clusters of NCCs based on the biochemistry and transport of molecular signals. On this more fundamental level, we first show that CIL and COA arise naturally from the underlying reaction and diffusion of Rho GTPases. Second and more importantly, we demonstrate that CIL and COA, acting on a randomization scheme for cell polarity, produce persistence of cell polarity, and consequently spontaneous collective migration of NCCs. This provides a plausible biological basis for POP from the biochemistry of Rac1 modulation by CIL and COA. As it turns out, POP is not an additional rule to be posed alongside CIL and COA. It is in fact a natural outcome of CIL and COA, as well as a conduit through which these two fundamental mechanisms give rise to the observed spontaneous collective migration.

## 2 Methods

Our model draws from a growing literature on computing cell migration from signaling proteins in the interior and on the membrane of cells [19, 20, e.g.]. Our general conceptualization of the NCC collective migration is as follows. Polarization and protrusion of individual cells are governed by Rho GTPases on the membrane [21–23], subject to turnover between the membrane-bound and cytoplasmic forms of the signaling proteins. In many cell types, Rac1 promotes F-actin assembly in lamellipodia at the protrusive front of the cell, while RhoA enhances myosin-induced cell contraction at the rear [24,25]. For NCCs, in particular, Rac1 and RhoA have been identified as the key proteins modulating CIL and COA [6,18]. Therefore, our model only accounts for Rac1 and RhoA and omits other GTPases such as Cdc42 and various downstream regulators. The level of active Rac1 and RhoA on the membrane determines the protrusive and contractile forces on the membrane and in turn the deformation and movement of the cell. Cell-cell interaction occurs through modulating each other’s Rac-Rho biochemistry. For example, cell-cell contact inhibits Rac1 and elevates RhoA at the site of contact in both cells. Thus, the cell protrusions retract and the cells move apart. Finally, the Rac-Rho dynamics is subject to a random noise so as to produce the tortuous trajectory commonly seen for single migrating cells.

To implement the above ideas, we represent each cell by a polygon of *N* vertices connected by elastic edges (Fig. 1). These edges represent the membrane-cortex complex [26], similar to the vertex models widely used for epithelial morphogenesis [27, 28]. Rac1 and RhoA levels are defined on the vertices as well as in the cytoplasm. A kind of “mesh refinement” has been tested, and balancing accuracy with computational cost, we have chosen *N* = 16 for all simulations in the rest of the paper. For details see Fig. S1 in the Supporting Information (SI).

**Figure 1:**
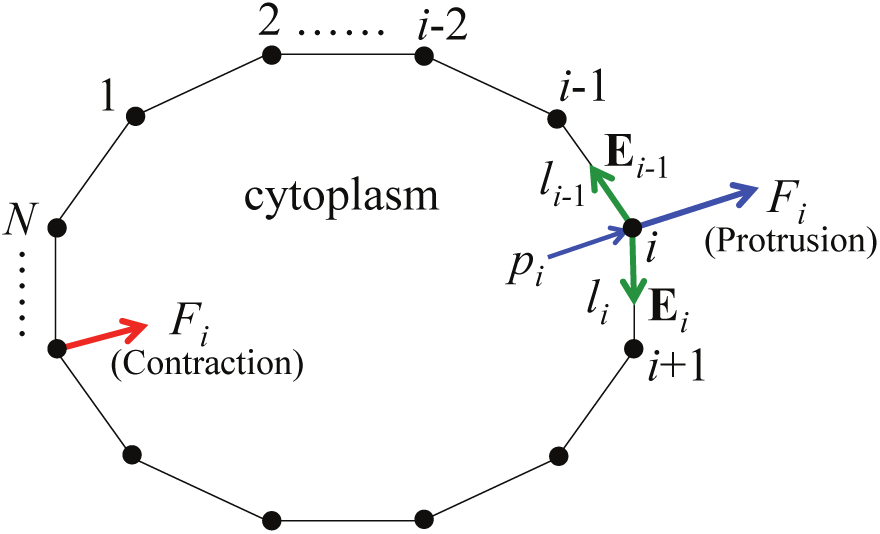
A model cell represented by a polygon of elastic edges. The vertices bear active and inactive levels of Rho GTPases and move according to the various forces acting on them. The edge between vertices *i* and *i* + 1 has length *l*_*i*_ and carries an elastic tension of **E***_i_*. The normal forces include pressure *p*_*i*_ and a protrusion or contraction force *F*_*i*_.

We imagine the cells being spread on a substrate, but do not explicitly account for focal adhesions. Each vertex on the cell membrane is subject to 2D forces acting in the plane: a pressure force from the cytoplasm enclosed by the cell membrane, cortical tension in the membrane segments, and a protrusion or contraction force determined by the Rac1 and RhoA levels on the vertex. As a result, the vertices move, without inertia, at a speed determined by the resultant force and a friction factor. The biochemical and mechanical components of the model are intimately coupled. For clarity of narration, however, we will describe each in turn.

### 2.1 Biochemistry of Rac1 and RhoA

For both Rac1 and RhoA, the model tracks three forms of the signaling protein: the active membrane-bound form, the inactive membrane-bound form, and the inactive cytosolic form. For Rac1, we normalize the amount of these forms by the total amount of Rac1 in the cell (details in SI), and denote them as *R*^*a*^, *R*^*i*^ and *R*^*c*^, respectively. Similarly, we denote the normalized amounts of RhoA by *ρ*^*a*^, *ρ*^*i*^ and *ρ*^*c*^. Note that the membrane-bound forms are defined on the cell boundary vertices, and may exhibit spatial distributions. In fact, cell polarization will be marked by spatially segregated distributions of active Rac1 and RhoA. The cytosolic levels are functions of time but not space. Since the bulk diffusion of Rho GTPases inside the cytosol is much faster than on the membrane, we assume the cytosol to be well mixed [19, 29, 30]. The total amounts of Rac1 and RhoA are each conserved.

The biochemistry of Rac1 and RhoA can be represented by a set of reaction-diffusion equations. We assume that on the membrane, the active and inactive forms of each protein interconvert with activation and deactivation rates, denoted by *K*^+^ and *K*^-^ for Rac1 and *κ*^+^ and *κ*^-^ for RhoA. Only the inactive form of protein may dissociate from the membrane to diffuse within the cytosol. The membrane association and dissociation rates are denoted by *M* ^+^ and *M*^-^ for Rac1, and by *μ*^+^ and *μ*^-^ for RhoA. Using Fick’s law to compute the 1D diffusion flux *J* of the active and inactive Rac on the membrane, we discretize the reaction-diffusion equations as follows (see SI for details):

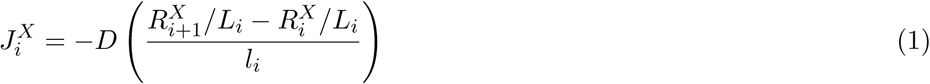

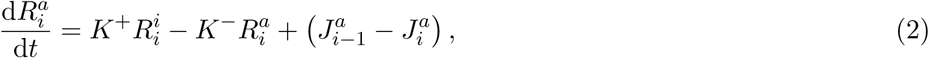

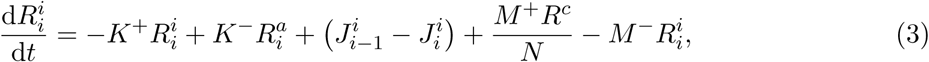

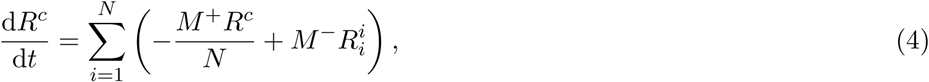

where the subscript *i* indicates the *i*^th^ vertex on the cell membrane, *l*_*i*_ is the edge length between vertex *i* and *i* + 1, *L*_*i*_ is the average of *l*_*i*_ and *l*_*i-1*_, *D* is the diffusivity on the membrane, and *N* is the total number of vertices. *J*_*i*_ approximates the diffusive flux from vertex *i* to vertex *i* + 1, its superscript *X* being *a* or *i* for the active and inactive forms of Rac.

RhoA obeys similar kinetic equations:

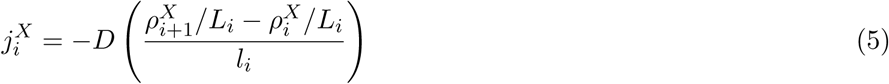

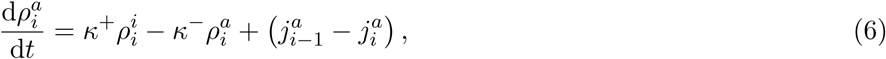

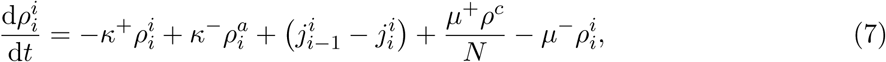

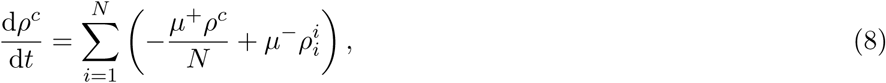

The equations imply conservation of the total amount of each Rho GTPase species.

These biochemical interactions are central to the model’s ability to reproduce cell polarization, stochastic repolarization (i.e. random changes in migration direction), CIL and COA, all of which are encoded in the Rho GTPase activation and inactivation rates *K*^*±*^ and *κ*^*±*^ (See SI for algebraic details). We design these rate functions according to biological observations and prior modeling in the literature. In the following we briefly discuss the modeling of each of these effects.

- *Polarization.* To capture cell polarity, the activation rates of Rac1 and RhoA each reflect the species’ autocatalytic capacity, while their de-activation rates reflect mutual inhibition on the cell membrane [30]. Following [29, 30], we represent the Rac and Rho auto-activation and mutual inhibition through Hill functions, which allow spontaneous polarization of cells with Rac1 and RhoA peaking on opposite sides of a cell. This polarity is the precursor of cell motility.
- *Stochastic repolarization.* Similar to other migratory cells [31], NCCs intrinsically produce random Rac1-mediated protrusions [11] that can out-compete existing protrusive fronts and change the cell’s existing polarized morphology. To capture this, every *T*_*r*_ minutes we randomly select a subset of vertices and up-regulate the Rac1 activation rate on them. Aside from the reaction and diffusion, competition between protrusions is also mediated by a negative feedback of the membrane-cortex tension on Rac1 activation. This is based on observations that as a new protrusion raises the membrane-cortex tension globally over a migratory cell, the elevated tension not only resists actin protrusions mechanically, but also through biochemical signaling that inhibits Rac1 [32] or the SCAR/WAVE2 complex downstream of Rac1 [33]. Thus, hotspots of Rac1 activity compete with each other on the membrane, allowing an upstart to replace an existing protrusion on occasion.
- *Contact inhibition of locomotion.* Contact between two NCCs is known to activate the noncanonical Wnt signalling pathway, which results in the down-regulation of Rac1 and the up-regulation of RhoA [18], leading to CIL. This is effected in the kinetic equations by CIL factors that elevate the Rac1 deactivation rate *K*^−^ and the RhoA activation rate *κ*^+^ on any vertex that has come sufficiently close to a neighboring cell. As a result, the polarity of two cells approaching each other is modified, with the protrusions retracting and the cells shrinking from each other.
- *Co-attraction.* Previous work has demonstrated that binding between the NCC autocrine C3a and its receptor C3a leads to up-regulation of Rac1 [6]. This enhances protrusion toward each other between neighbors and produces COA. Since C3a diffuses through the extracellular matrix at much faster timescales than NCC migration, we need not model the diffusion of C3a explicitly, but rather assume a steady state exponential distribution of C3a surrounding each NCC [13]. In our model, we realize COA by up-regulating the Rac1 activation rate on any vertex of a cell that is sufficiently close to a neighboring cell to “sense” the C3a distribution of the latter.

### 2.2 Mechanics of cell deformation and motility

The cell being represented as a polygon (Fig. 1), its shape and movement is specified by the position **r** of each vertex and its speed 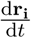.Thus, we write

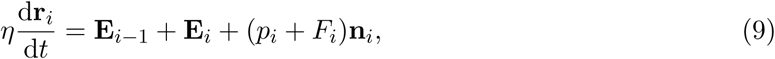

where **E***_i_* is the elastic tension along the edge between vertices *i* and *i* + 1, *p* is the cytoplasmic pressure, *F* is the protrusion force due to actin filaments on the membrane, and **n** is the unit outward normal vector at vertex *i* (Fig. 1). The elastic tension **E** is proportional to the strain in each edge, relative to an undeformed length of *l*_0_, with an elastic modulus *λ*. The cytoplasmic pressure is such that a reduction in the cell area is resisted by an outward normal force:

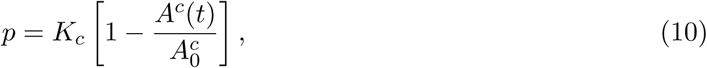

where *A*^*c*^(*t*) is the cell area at time *t*, *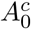* is its resting area, and *K*_*c*_ is the cytoplasmic stiffness. Expansion of the cell area is constrained by the membrane elasticity *λ*. As our model is 2D, conserving the cell area is the counterpart of conserving cell volume in 3D. We may think of *A* as the cell area viewed from above if we take the height of the cell to be constant.

The active protrusion or contraction force is determined by the activated levels of Rho GTPase at the vertex. If the active Rac1 is greater than active RhoA, the active force is protrusive (positive). Otherwise it is contractile (negative):

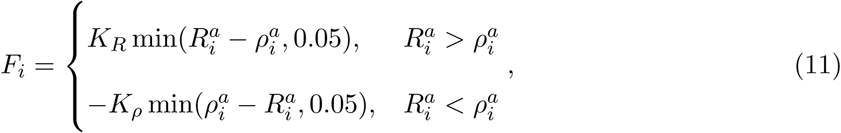

where *K*_*R*_ and *K*_*ρ*_ are constants governing the magnitude of the Rac1 and RhoA forces, respectively. In both cases, the force is capped at a constant maximum magnitude when the difference in the GTPase levels exceeds 0.05. The rationale for this functional form is explained in SI.

The model has a list of geometric, physical and kinetic parameters. These are tabulated in the SI, along with the sources from which we have determined their values for use in the following simulations.

## 3 Results

### 3.1 Single-cell behavior: polarization and random walk

The first and most basic prediction of the model is that a single cell spontaneously polarizes and migrates (Fig. 2; also see Movie 1 in SI). As initial condition for *t* = 0, we divide the total amount of Rac1 into 10% membrane-bound active, 10% membrane-bound inactive and 80% cytosolic. The membrane-bound active and inactive Rac1 is each randomly distributed among the membrane nodes. A similar scheme is followed for RhoA. Unless explicitly stated otherwise, such a random initial condition is used in all subsequent simulations, including multiple realizations from different random initial conditions for gathering statistical information.

**Figure 2:**
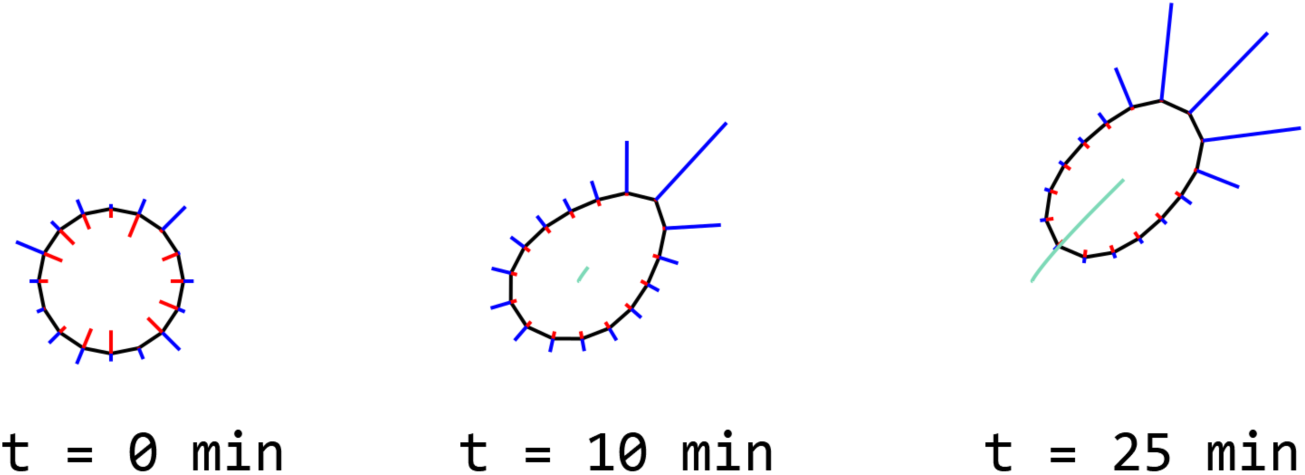
A single cell (diameter *d* = 40 μm) polarizes and migrates. The lengths of line segments pointing in or out are proportional to the local levels of active GTPases (outwards blue lines for Rac1, inwards red lines for RhoA). The distance of migration is indicated by the trace of the cells centroid.

The polarization is realized via the wave-pinning mechanism described by Edelstein-Keshet and coworkers [29, 34]. In essence, the nonlinear autocatalysis of Rac1 or RhoA coupled with a finite total amount of the proteins ensures the coexistence of multiple solutions corresponding to a lowor high-activity state. The mutual inhibition between Rac1 and RhoA leads to complementary distributions of these two species, with one end of the cell featuring high Rac1 and low RhoA and the opposite end low Rac1 and high RhoA. The former becomes a protruding front and the latter a retracting rear. Thus the cell moves forward. Note that cell-substrate adhesion is not explicitly accounted for, nor are stress fibers. The steady-state migration speed has been tuned to the experimentally observed 3 μm min*^-1^* by adjusting the coefficient *K*_*ρ*_ for the contractile force (Eq. 11) relative to the protrusive force and friction (Eq. 9). See Table 2 of SI for details.

The model also captures random repolarization and changes in the direction of motion. As a result, the trajectory of the single cell resembles that of a random walker (Fig. 3*a*). As explained in Subsection 2.1, the random change in polarization is realized in our model by periodically upregulating the Rac activation rate *K*^+^ at randomly selected vertices. Thus, these vertices become potential hotspots of Rac1 activity. They compete with existing Rac1 peaks through corticaltension-based inhibition [32] and the polarizing nature of Rac1-RhoA chemical dynamics [29]. If a new hotspot supersedes an existing one and becomes a new protrusion, the cell repolarizes and changes its direction of migration (see Movie 1).

**Figure 3:**
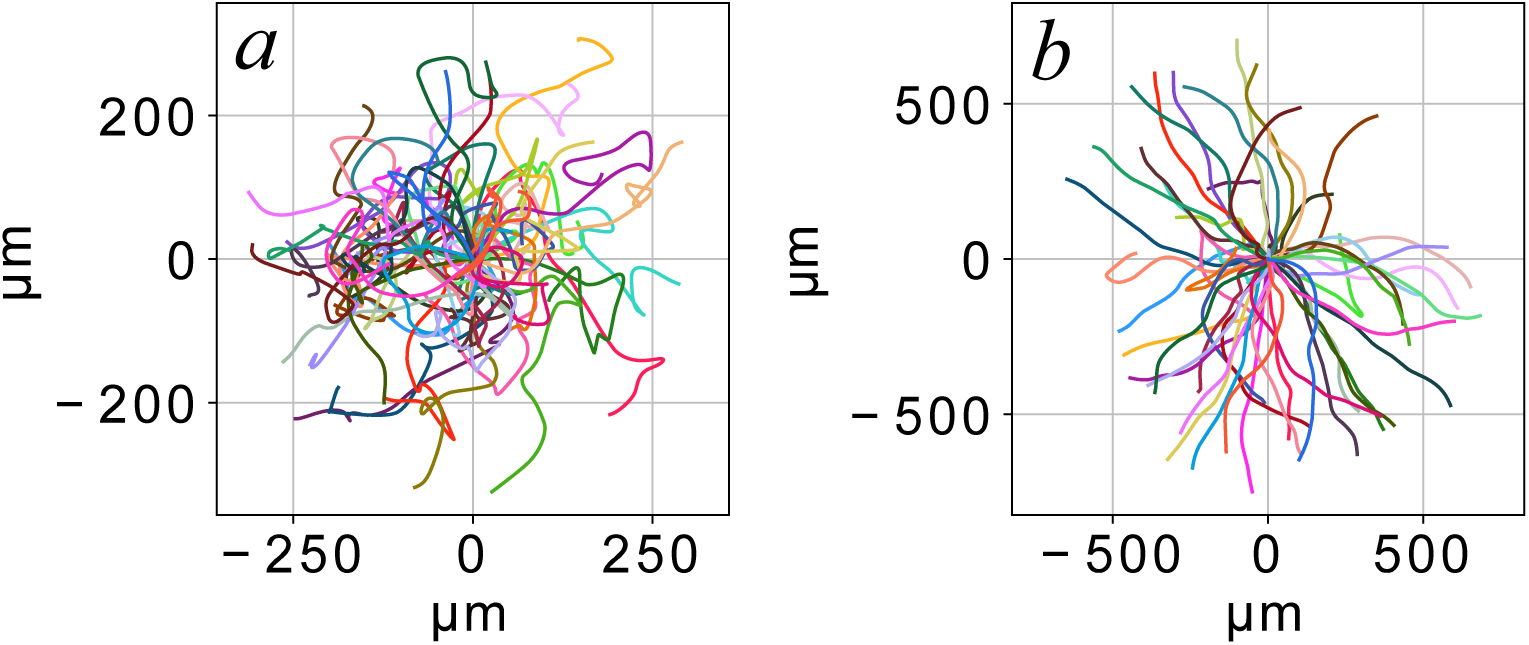
(*a*) Fifty single-cell trajectories show considerable tortuosity due to the periodic repolarization scheme in our model. Each trajectory starts from the origin with a randomly assigned initial Rac and Rho distribution and lasts 4 hours. The persistence ratio is *R*_*p*_ = 0.4 *±* 0.17 and the persistence time *T*_*p*_ = 26 *±* 24 min. (*b*) Increased persistence in cell migration when the Rac1 baseline activation rate 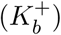 is reduced to 40% of its usual level. *R*_*p*_ = 0.85 *±* 0.17, *T*_*p*_ = 725 *±* 1140 min.

We have included random repolarization in our model since such changes in migratory direction are a common feature of single-cell migration for several cell types [35, 36]. As will be discussed below, it is also essential for capturing the spontaneous collective migration of a cluster of cells. *In vitro*, mammary epithelial cells alternate between two modes of migration: a highly directional phase of cell motion, and a re-orientation phase in which the cell produces new leading edges and sharply changes its direction of motion [37]. This corresponds closely with our model predictions. Theveneau *et al.* [38] reported similar bimodal behaviour of a single neural crest cell (see their Supplementary Information, Movie S5), and each phase lasts between 15 and 20 min. In our model prediction, the directional motion and repolarization each takes about 20 minutes. These periods are determined by the kinetic rates and the parameter *T*_*r*_, the time period between each new choice of random up-regulation of Rac1.

One way to quantify the random changes in migratory direction is through the persistence ratio *R*_*p*_, defined as the end-to-end distance of a cell’s trajectory divided by the path length of the tortuous trajectory [14, 37, 39]. Since the cell essentially executes a random walk, the end-to-end distance increases with time as 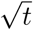 while the contour length as *t*. Thus, 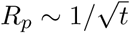 will in generally decrease with *t*, and is meaningful only if measured for a specific time period. Figure 3(*a*) depicts multiple trajectories of a single cell generated from different random initial conditions in the model. The average persistence ratio is *R*_*p*_ = 0.4 over a period of 4 hours. For single neural crest cells, *in vitro* experimental data [11, 40, 41] suggest *R*_*p*_ values ranging from 0.1 to 0.4 over unspecified period of time, making comparison difficult.

A more general quantitative measure of the randomness of single-cell motility is the persistence time *T*_*p*_ [39], defined as the characteristic time for the decay of the directional autocorrelation [42]. Unlike the persistence ratio *R*_*p*_, *T*_*p*_ is independent of the observation period and thus more convenient for comparisons. The average *T*_*p*_ is around 26 minutes in our simulation (Fig. 3*a*). We did not find any experimental data of *T*_*p*_ for NCCs, but human mammary epithelial cells have a persistence time of *T*_*p*_ *∼* 10 min [37] while fibroblasts exhibit a range of persistence times between 15 to 120 minutes [43, 44].

Pankov *et al.*’s experiment [31] showed that inhibiting Rac1 GEFs (activators) using a drug led to increased cell persistence, because cells produced weaker Rac1 hotspots to compete with established ones. Our model has captured this feature. Figure 3(*b*) shows cell trajectories predicted by the model using the same randomization scheme, but with the baseline rate of Rac1 activation reduced to 40% of its normal value to model the effect of a drug that globally inhibits Rac1 activation. This drastically increases cell persistence from 0.4 to 0.85, and the persistence time from 26 to 725 min. The cell almost moves in straight lines in this case, since random spikes in Rac1 activation are too weak to override the cell’s existing polarity. This feature will be relevant to our modeling of persistence of polarity as discussed in Subsection 3.4 below.

### 3.2 Contact inhibition of locomotion

Scarpa *et al.* [41] carried out *in vitro* experiments to study CIL between two cells placed on a fibronectin strip. Thus, the cells are confined to effectively 1D movement in a corridor, the area outside being non-adhesive and prohibitive for the cells. To simulate this experiment, we place two cells in a corridor with boundaries designed to be repulsive to the cell via the same biochemical pathways as underlie CIL. That is, the boundary will up-regulate the RhoA activation rate as well as the Rac1 deactivation rate (Eqs. S4, S9 in SI). As an initial perturbation, we raise the Rac1 level on the parts of the cells facing each other. This gives them a polarity such that they initially move toward each other. We also turn off COA in this subsection for a cleaner manifestation of CIL. The sequence of the simulation is illustrated in Fig. 4 and Movie 2.

**Figure 4:**
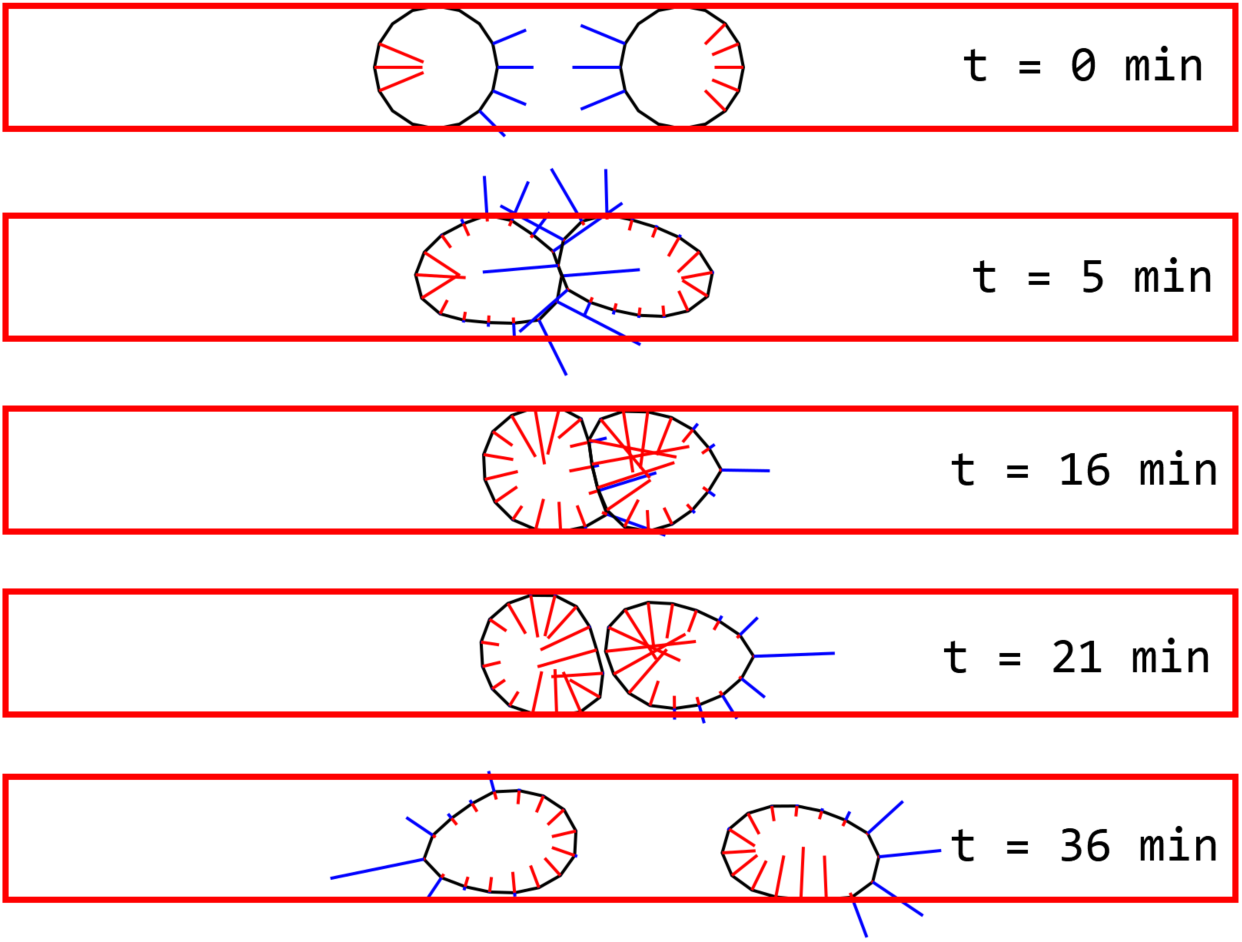
A typical CIL interaction in a corridor of one-cell-diameter width. Contact between the cells causes local activation of RhoA, which in turn reverses the cell polarization due to Rac1-RhoA chemical dynamics. See also Movie 2 in SI.

Upon contact, the vertices at the cell’s front receive a boost in their rates of RhoA activation and Rac1 deactivation. This quickly extinguishes the Rac-rich protrusion fronts (at 16 min) and subsequently turns them into Rho-rich retraction fronts (21 min). As the result of reaction-diffusion and global conservation of the GTPases (e.g., Eqs. 2–4), there is now an abundance of inactive Rac1 on the cell membrane and a dearth of active RhoA away from the contact area. Besides, the retraction relieves the membrane tension and its inhibition of active Rac1. These favorable conditions promote the appearance and growth of tentative Rac hotspots on the opposing end of each cell (21 min). In time, these hotspots develop into new protrusion fronts pointing away from the regions of cell-cell contact, and the two cells move apart (31 min). The sequence of contact-inhibition agrees closely with the experimental observations of Scarpa *et al.* [41] (cf. their Fig. 3c,d and Movie 1).

### 3.3 Co-attraction

To demonstrate the effect of COA, we arrange 49 cells in a 7 by 7 initial square (Fig. 5*a*), each bearing a random initial distribution of Rac1 and RhoA as described above for a single cell. The cells spontaneously polarize and move, as demonstrated in Fig. 2 for single cells. Meanwhile they interact through CIL. Here we compare two simulations, with COA turned on and off. See also Movies 3(*a*,*b*) in the SI.

**Figure 5:**
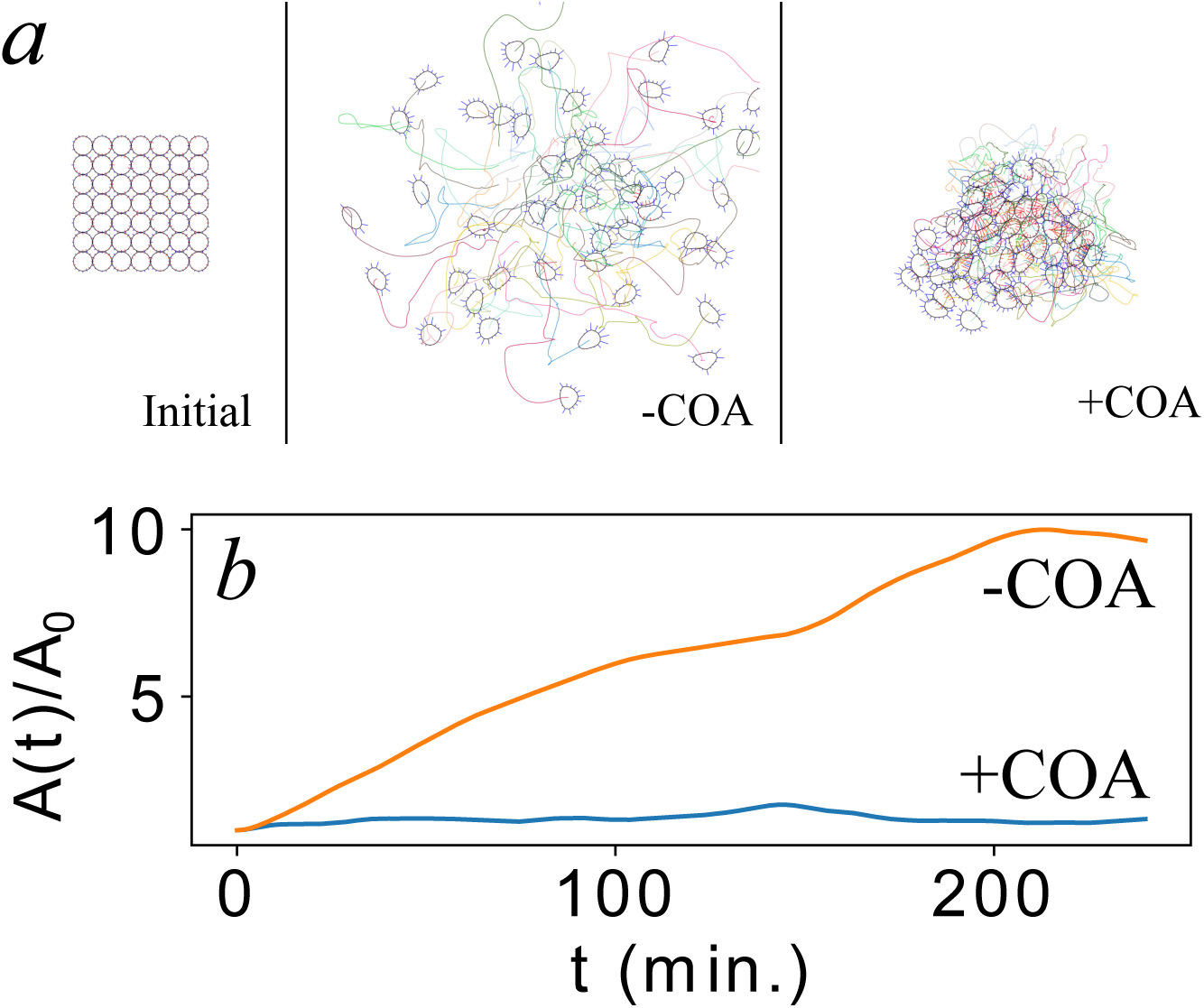
The effect of COA for a cluster of 49 cells. (*a*) Starting from a square initial array, 49 cells scatter in 4 hours with COA turned off (-COA) but stays in a cluster with COA turned on (+COA). (*b*) Temporal evolution of the area of the cell cluster *A*(*t*) normalized by the initial area *A*_0_, with and without COA.

With COA turned off, we see that the cells scatter as a result of their random depolarization and CIL (Fig. 5*a*). The total area of the cluster *A*, defined as the area of the convex hull of the cell centroids, increases roughly linearly in time (Fig. 5*b*). This resembles the scattering of Brownian particles. With COA turned on, however, the cluster retains its integrity in time (Fig. 5*a*). Its total area fluctuates around a mean value: *A/A*_0_ = 1.5 *±* 0.18 (Fig. 5*b*). This illustrates how COA, modeled on the activation of Rac1 through the C3a/C3aR ligand-receptor pathway, helps keep NCCs clustered and thereby allows for continued intercellular contact and interaction. This will be shown next to be essential for a group’s spontaneous collective migration.

### 3.4 Persistence of polarization

As explained in the Introduction, analyzing prior models and experiments has led us to believe that a degree of persistence of polarity (POP) is necessary for spontaneous collective migration of neural crest cells. To illustrate this persistence and to explore its origin, we will first analyze the two-cell migration simulation depicted in Fig. 6 and Movie 4.

**Figure 6:**
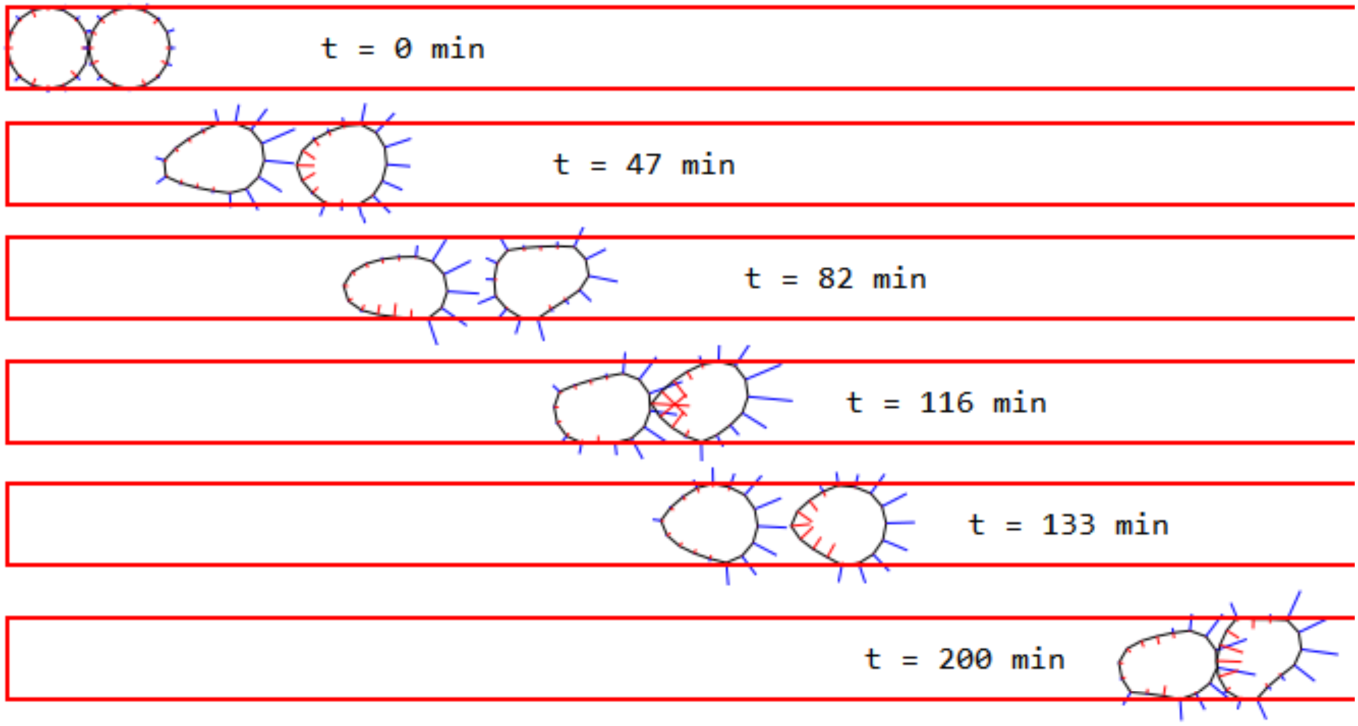
Persistence of polarity exhibited by two cells migrating within a 1-cell-diameter wide corridor. Highly persistent group motion develops due to the trailing cell constantly neutralizing any new protrusions on the rear end of the leading cell through CIL, and the leading cell reinforcing forward protrusions on the trailing cell through the effect of COA.

At the start of the simulation, the two cells are placed next to the left end of the corridor, whose boundaries confine the cells via CIL as explained in connection to Fig. 4. Because of this confinement and the initial asymmetry in the geometry, the two cells develop protrusion fronts toward the right, and the cells start to move to the right (*t* = 47 min). What is somewhat surprising is that this motion is sustained for the entire duration of the simulation (10 hours) across some 40 cell diameters, the initial portion being shown in Fig. 6. This is in contrast to the behavior of a single cell, which, after the initial geometry-induced asymmetric motion away from the left end, essentially adopts a 1D random walk in the corridor with no persistent net movement in either direction.

Comparing the persistent two-cell motion with the random walk of the single cell, we first note that COA keeps the two cells within close proximity through the entire duration. This ensures the continual interaction between the two through CIL and COA. The effect of CIL on the leading cell is such that it never develops a viable new protrusion in the rear. Any such hotspot, as appears at 82 min, is quickly extinguished by CIL as the budding protrusion comes into contact with the trailing cell (116 min). If such contact weakens the protrusion front of the trailing cell, this effect is short-lived as a slow-down of the trailer will end the contact. The nascent Rac1 hotspot on the rear of the trailing cell, visible at 133 min, cannot compete with the forward protrusion that is reinforced by COA. Thus, the pair continues its directional migration, and similar cycles of interaction repeat in time (e.g. 200 min). Throughout the entire migration, forward protrusions are long-lived and dominant on both cells, while rear protrusions are ephemeral and immaterial. This is clearly seen from the distribution of the life-time of protrusions in different directions (Fig. 7). The persistence ratio for the two-cell migration is *R*_*p*_ = 0.98 over 10 hours, while for a single cell *R*_*p*_ = 0.23 over the same period. Also the persistence time is *T*_*p*_ = 443 min for the pair and only 20 min for the single cell. Therefore, COA and CIL act together to suppress the bursts of Rac1 up-regulation on the cell membrane, which would have produced random repolarization on a single cell. Persistence of polarity (POP) arises naturally from the cooperation between COA and CIL, and perpetuates the initial asymmetric motion of the cells endowed by the geometric confinement. This is the most important insight gained from this study.

**Figure 7:**
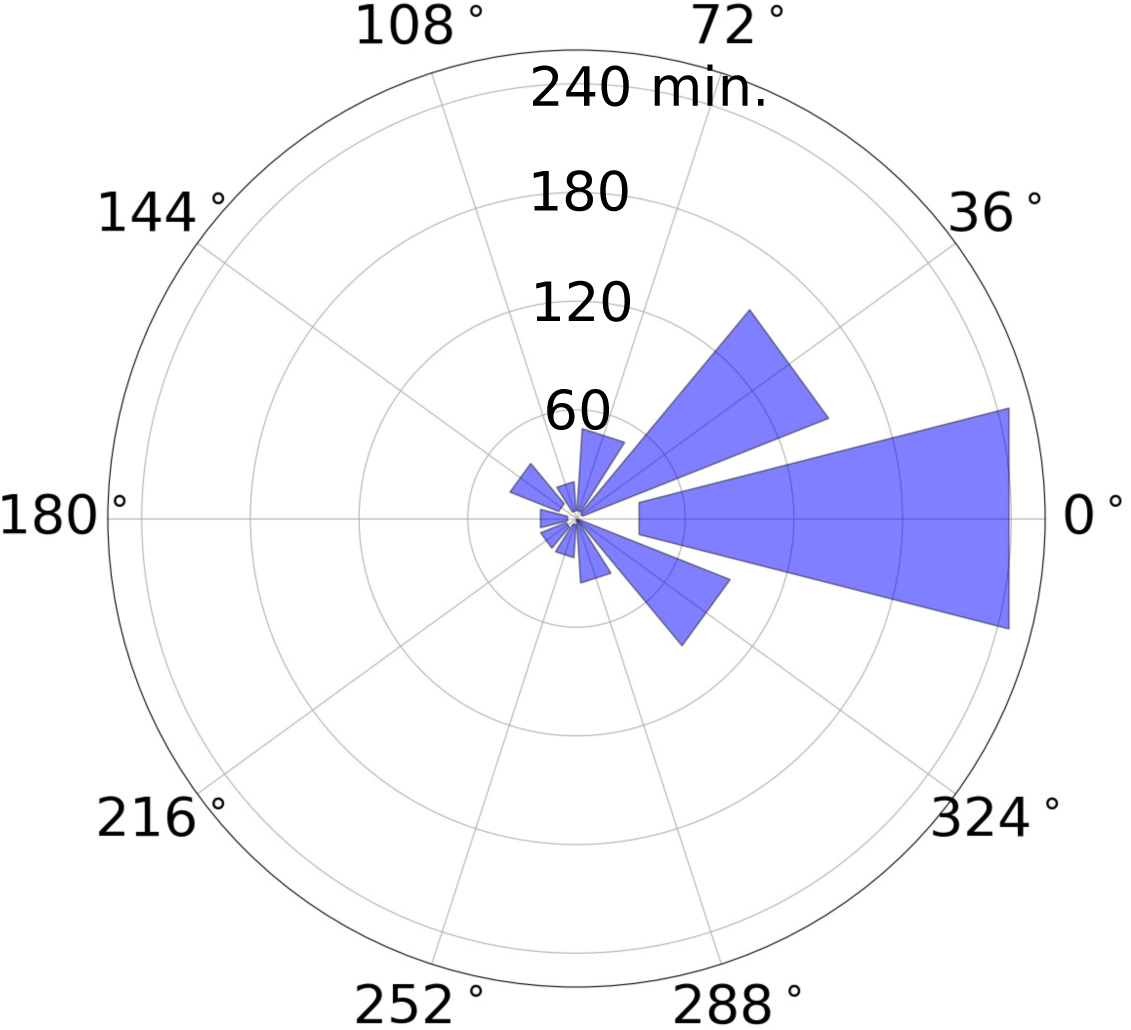
Distribution of the lifetime of protrusions shows a strong bias in favor of forward protrusion down the corridor. We define a protrusion as a vertex on which the active Rac1 is at a level of at least 25% of the maximum among all vertices, and exceeds the active RhoA. Dividing 2*π* into 10 intervals, we collect the lifetime of protrusions oriented in each angular interval on the two cells of Fig. 6 throughout the simulation. The sector centered in each interval is a Tukey box marking the first and third quartiles of the spread.

This insight answers the main questions that had motivated our model. If CIL and COA are postulated as ad hoc rules on the *supracellular* scale, they are insufficient for spontaneous collective migration. From this we have hypothesized that POP is a necessary third ingredient. Interestingly, having built our model from the underlying GTPase dynamics on the *intracellular* scale, we find that POP emerges from the collaboration between COA and CIL, and need not be added separately after all. What has been added to our model, and missing from prior rule-based models [13, 14], is the Rac-Rho biochemistry that allows a description and rational explanation of random repolarization. By stunting new Rac1 hotspots and suppressing this random repolarization, COA and CIL gives rise to POP. In an earlier version of our model, we realized repolarization by periodically erasing the existing polarization of a cell and imposing a random new Rac1 and RhoA distribution. This did not allow Rac1 suppression by CIL and COA, and failed to yield POP (Fig. S2 and Movie 5).

The idea that Rac1 suppression promotes persistence in single-cell migration is well established in the experimental literature. Pankov *et al.* [31] showed that human fibroblasts with Rac1 activity suppressed by RNA interference or the Rac GEF inhibitor NSC 23766 had increased cell persistence. This was due to a decrease in the strength and number of random Rac1-mediated protrusions forming on the cell periphery that could otherwise become dominant and re-orient the cell’s polarity. Later, Bass *et al.* [45] observed that the binding of membrane protein Syndecan-4 with fibronectin suppressed Rac1 to produce highly persistent fibroblast migration. Syndecan-4-null fibroblasts migrated randomly with delocalized Rac1 activity. Subsequently, Matthews *et al.* [46] showed that the same Syndecan-4 mediates increased persistence of migration in NCCs. As noted in connection to Fig. 3(*b*), reducing the Rac activation rate globally does markedly increase the persistence in the motility of a single cell in our model. Qualitatively, this corresponds to Rac suppression by drugs [31] or Syndecan-4 [45, 46]. In our simulation of a pair of interacting NCCs (Fig. 6), COA ensures continual action of CIL, which reduces Rac1 activation. Thus, the simultaneous action of COA and CIL amounts to a pathway to POP that resembles the GEF inhibitor or Syndecan-4 in single-cell migration. Turning off either COA or CIL compromises POP in our model (Fig. S3.)

Finally, we should note that POP is stochastic in nature and is not foolproof. If, by chance, new Rac1 hotspots appear simultaneously on the rear end of both cells, they may overcome the forward polarity and cause the pair to reverse course. Whether this occurs depends on the initial Rac1 and RhoA distributions, and on the number of cells in the cluster. We will discuss the robustness of POP at greater length in the next subsection.

### 3.5 Spontaneous collection migration of larger clusters

The key features of the persistent migration of two cells carry over to the collective migration of clusters comprising more cells. Figure 8 and Movie 6 demonstrate the spontaneous collective migration of 49 cells down a corridor. The cluster migrates persistently as a whole for about 20 cell diameters during a 10-hour period. Although the cluster area fluctuates, COA keeps the coherence of the group so that the mean area does not increase over time.

**Figure 8:**
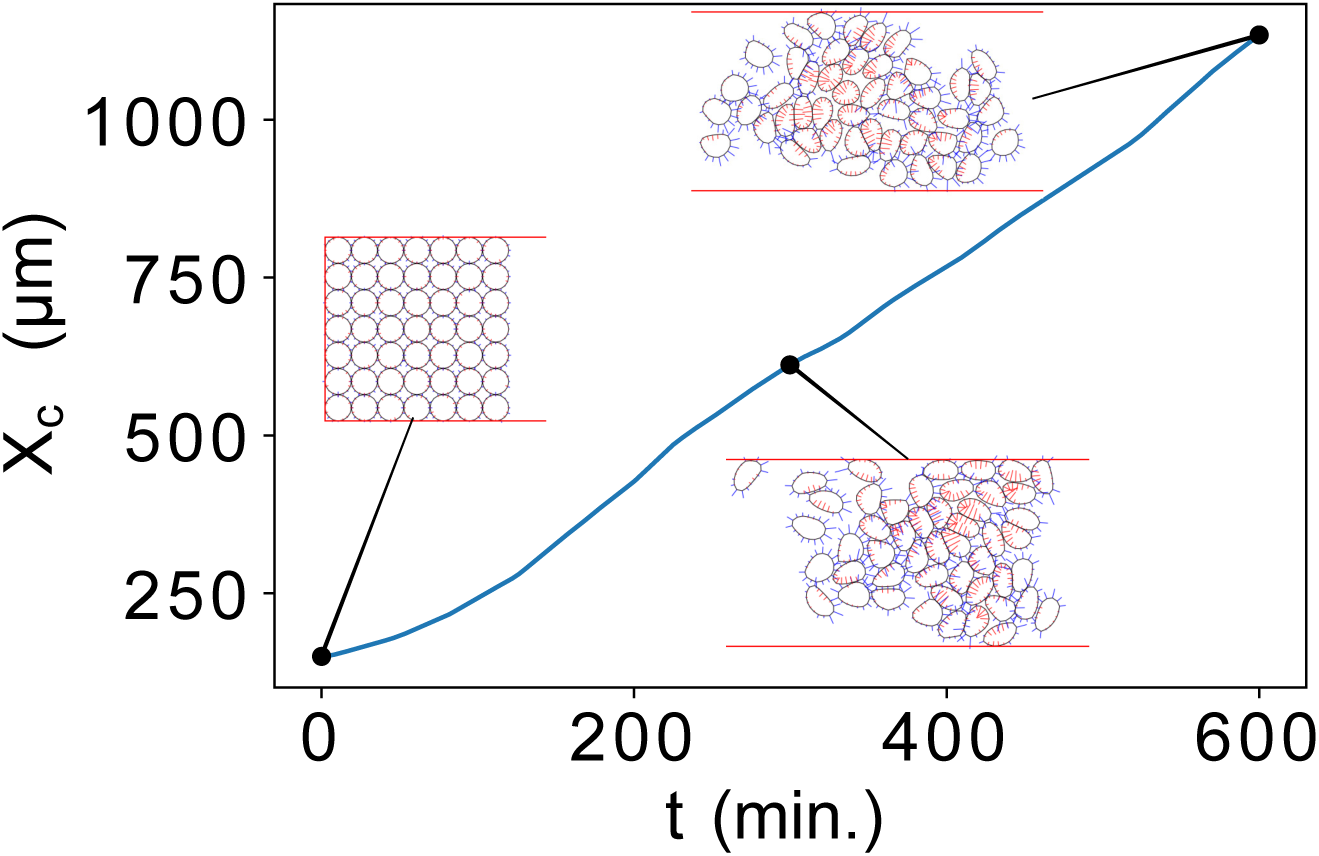
Spontaneous collective migration of a 49-cell cluster down a corridor of width *w* = 7. *X*_*c*_ denotes the cluster centroid location and the snapshots show that the cluster retains its integrity while migrating collectively. See Movie 6 in SI for a more detailed view of the migration.

To quantify the cluster size effect on the spontaneous collective migration, we have to distinguish it from the confinement effect of the corridor. With a larger number of cells inside the same corridor, the cluster will be more crowded and this amounts to an effectively stronger confinement. As an approximate way of separating the two effects, we adopt the following scheme in the next two subsections. In this subsection, we keep the width of the corridor *w*, scaled by the cell diameter *d*, as 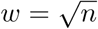, *n* being the total number of cells. Thus, we always have a square initial array of cells enclosed on three sides by walls, as in Fig. 8, *t* = 0. This gives us, in a sense, “equal confinement” among the different *n*. In the next subsection, we will vary *w* for a fixed *n* to examine the effect of confinement.

Another complication is that in our model for COA, the overall level of C3a is the sum of all such signals emitted from each cell. Thus, a larger number of cells raise the level of C3a around them. As a result, we do not need as strong a COA intensity per cell for larger *n*. By modulating the maximum COA strength *M*_COA_ (See Eq. S3 of SI for definition), we ensure that with increasing *n*, the average separation *S* between neighboring cells remains roughly the same. This is illustrated in Fig. 9(*a*). *S* is calculated by taking a Delaunay triangulation of the cell centroids and averaging all its edges, and then time-averaging this separation over the duration of the simulation. For each *n*, we have run 20 independent realizations of the simulation starting from random initial Rac1 and RhoA distributions. The error bars indicate the spread among the 20 runs. Going from *n* = 4 to 49, we have reduced *M*_COA_ from 24 to 8. These values are tabulated in SI.

**Figure 9:**
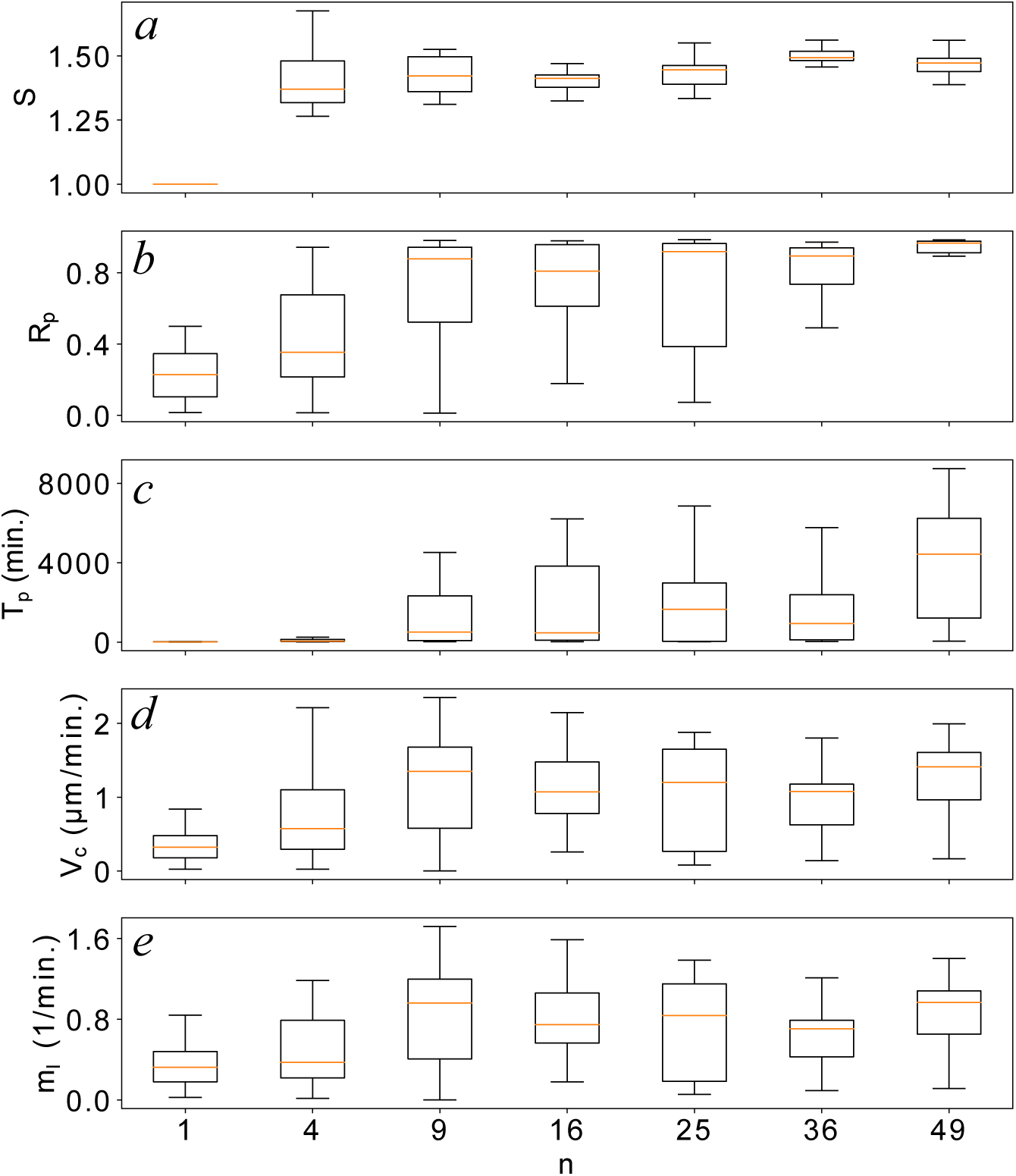
Effect of the cluster size *n* on key features of spontaneous collective migration. For each *n* value, the Tukey box-whisker represents 20 independent runs from different random initial GTPase distributions. The top and bottom of the box mark the first and third quartiles of the data, with the median indicated by the bar inside. The top and bottom of the whiskers mark 1.5 times the interquartile range from the edges of the box. (*a*) As *n* increases from 4 to 49, we have decreased the strength of COA to maintain roughly the same nearest neighbour separation *S* throughout the simulation. (*b*) The group persistence ratio *R*_*p*_ over 10 hours, computed from the trajectory of the centroid of the cluster. (*c*) The persistence time *T*_*p*_ of the centroid of the cluster. (*d*) The average migration speed *V*_*c*_ of the centroid of the cluster. (*e*) The migration intensity *m*_*I*_ = *V*_*c*_/*S*.

Figure 9(*b*–*e*) shows that the intensity of persistent collective migration increases with increasing *n*, rapidly for small clusters (*n* = 4 to 9) and mildly for the larger ones. Take the persistence time *T*_*p*_ in panel (*c*) for example. For *n* = 4, *T*_*p*_ averages among all runs to about 120 min. For *n* = 9, this has increased to 1627 min. Further increasing *n* lengthens *T*_*p*_ relatively modestly. The same trend is seen for the persistence ratio *R*_*p*_ (panel *b*) and cluster migration speed *V*_*c*_ (panel *d*). The bottom panel (*e*) depicts the “collective migration intensity” *m*_*I*_ = *V*_*c*_/*S*, which also increases with *n*. This ratio gives us a characteristic frequency at which the cells would pass a fixed location in the corridor. It is similar to the “transport ratio” defined for the percentage of cells passing a certain position in the corridor [14]. But the migration intensity does not depend on choosing an arbitrary “observation post”, and thus provides a more universal measure of the efficiency of collective migration.

The explanation for Fig. 9 lies in the fact that for small clusters, POP has a greater probability of failure, when the cluster reverses course as a whole. For larger clusters, the chance of POP failure is much reduced. Figure 10 illustrates the stochastic nature of POP fallibility. Take *n* = 9 for example. Among the 20 trajectories, the 10 that have traveled the farthest have never suffered a reversal, i.e. *V*_*c*_ *>* 0 for the entire duration. For the rest, reversal may first set in as early as *t* = 60 min or as late as 520 min. The reversal is invariably accompanied by a large portion of the cells simultaneously developing rear-facing Rac1 hotspots that survive and mature into protrusions. Fig. S4 and Movie 7 in the SI illustrate this process in detail for a 4-cell cluster.

**Figure 10:**
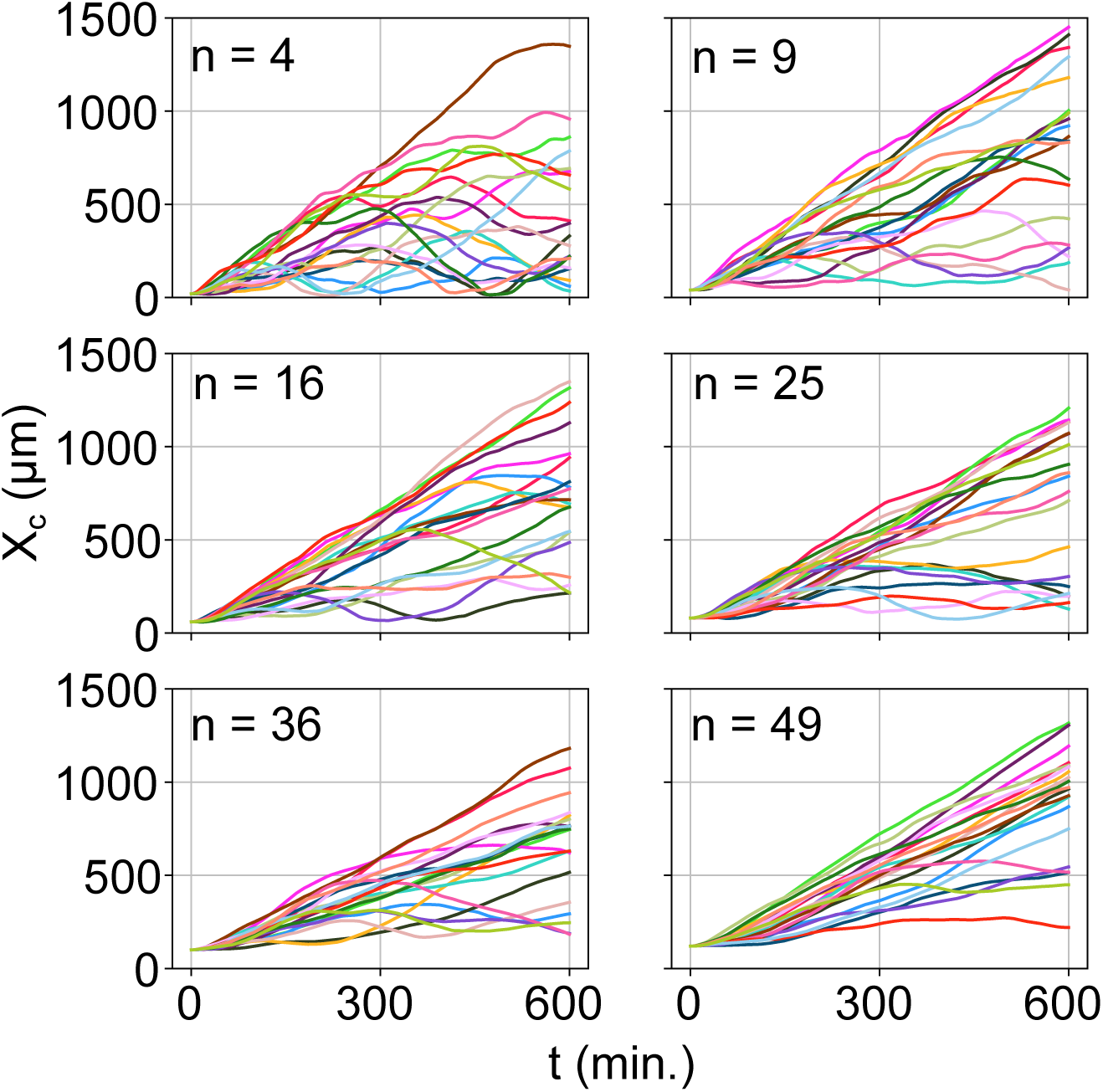
The trajectories of the cluster centroid *X*_*c*_(*t*) for various group sizes. The cell diameter *d* = 40 μm. Each plot includes 20 runs starting from different random initial Rac1 and RhoA distributions under otherwise identical conditions. Some of the trajectories suffer from reversals, but the collective migration becomes more robust with increasing *n*.

With increasing cluster size *n*, the probability of POP failure decreases, and the reversal tends to occur at later times if at all. As a convenient way to quantify the POP failure rate, we count the percentage of trajectories that end with a downward slope at the end of the 10-hour simulation. For *n* = 4, 50% of the 20 trajectories fail by this criterion. That percentage declines to 35% for *n* = 9, 25% for *n* = 16, 15% for *n* = 25, 20% for *n* = 36, and 10% for *n* = 49. Even though 20 realizations may not be a large enough sample size, the available results show a clear trend of more robust POP with increasing *n*. The failure of POP requires a large portion of the group to develop rear protrusions *simultaneously*. If only one or a few cells acquire rear hotspots among many, they will be quick extinguished by their neighbors through CIL. This explains why the probability of POP failure declines with increasing *n*. After all, as POP arises from cell-cell interactions, it is perhaps little surprise that it is more robust in larger clusters.

Finally, we can make some quantitative comparisons between our model prediction and experimental data. For NCCs migrating *in vivo* in *Xenopus laevis* embryos, Szabo *et al.* [14] reported a collective migration speed of *V*_*c*_ *≈* 1.5 μm/min. *In vitro*, the NCCs migrate at *V*_c_ *≈* 1 μm/min down a corridor coated with fibronectin and confined by borders that are rich in versican, an extracellular protein that repels NCCs. With the model parameters used (tabulated in SI), our model predicts an average speed *V*_*c*_ = 1.12 μm/min for the larger clusters (*n ≥* 9) (Fig. 9*d*), comparable with the experimental data. Note that in our model, we have adjusted the contractile force factor *K*_*ρ*_ (Eq. 11) to produce the experimental speed of 3 μm/min for a single cell in the rectilinear phase of motility. The agreement in cluster migration speed *V*_*c*_ is not the result of parameter fitting. We can also compare the persistence of the migrating clusters with experimental measurements. Szabo *et al.* [14] documented a persistence ratio *R*_*p*_ *≈* 0.85 over a time period of 4 hours *in vivo*, and *R*_*p*_ *≈* 0.87 *in vitro*. In Fig. 9(*b*), we have predicted an average persistence of *R*_*p*_ = 0.77 for the larger clusters (*n ≥* 9) over 10 hours. A caveat about comparison with experiment is that the model has a large number of parameters, not all of which can be ascertained directly from experimental measurements. The SI summarizes these parameters and their evaluation.

### 3.6 Confinement effect

In the corridor-based studies of Mayor *et al.* [6, 13, 14, 41] as well as in our simulations, confinement of the boundaries plays two important roles: to limit the migration to effectively one dimension, and to provide the initial geometric asymmetry that breaks the symmetry and ensures persistent migration later. Because of similar confinement effects *in vivo*, e.g. by versican-rich boundaries delimiting the migration of *Xenopus* cephalic NCCs, Szabo *et al.* [14] have investigated the role of confinement in NCC collective migration. Now we can examine confinement effects in our model as well.

For a fixed cluster size, *n* = 16, we have tested the efficiency of spontaneous migration in corridors of varying width (Fig. 11). At the start, the cells are arranged in a roughly rectangular shape that is bounded by walls on three sides at the left end of the corridor (Fig. S5). In Fig. 11(*a*), *w* = 1 appears to be a special case with much larger average neighbor separation than in the wider corridors. An examination of the cell trajectories shows that the 16 cells typically break up into two or more smaller groups that tend to drift apart (Fig. S6). This highly confined situation only allows longitudinal cell-cell interactions, which proves insufficient to maintain the single-file cluster as a whole. For this reason *w* = 1 cannot be compared with the wider corridors in Fig. 11.

**Figure 11:**
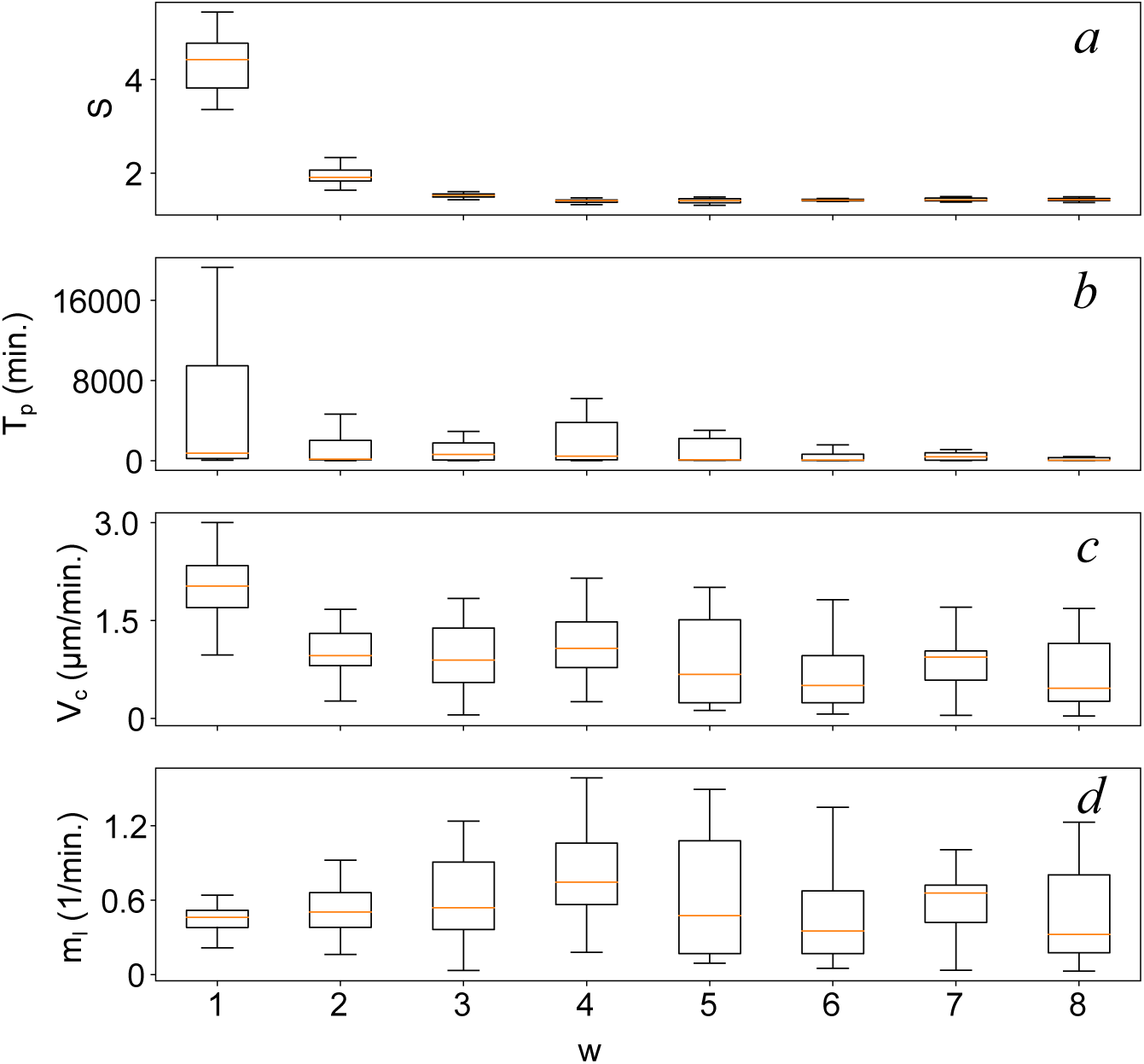
Confinement effect on the spontaneous collective migration of a cluster of 16 cells in a corridor of width *w*. In each panel the Tukey boxes represent 20 runs as in Fig. 9. (*a*) The average neighbour separation *S*. (*b*) The persistence time *T*_*p*_ of the centroid of the cluster. (*c*) The average migration speed *V*_*c*_ of the centroid of the cluster. (*d*) The migration intensity *m*_*I*_ = *V*_*c*_/*S*.

If we disregard the data for *w* = 1, then the most important feature of Fig. 11 is the existence of an optimal confinement, at 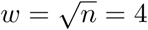 in this case, that produces the most efficient collective migration. This can be appreciated from each of the panels of the plot. For example, the average cell separation *S* is smallest for *w* = 4. In narrower corridors with 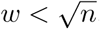, the strong confinement forces the cluster into an elongated array, and hampers the coordination between its front and rear via COA. Figure 12 illustrates this for *w* = 2. For 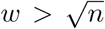, on the other hand, there are empty spaces between the top and bottom walls and the cluster (Fig. 12, *w* = 8). The weak confinement allows the cluster to spread along the *y*-direction, thus also enlarging *S*. Similarly, the cluster persistence time *T*_*p*_, cluster migration speed *V*_*c*_ and migration intensity *m*_*I*_ all attain maximum values at the optimal width. Generally, over-confinement in narrow corridors hampers COA and weakens coordination throughout the elongated cluster. Under-confinement in wide corridors allows the cluster to meander in a 2D plane instead of migrating down a 1D corridor. The width 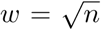 produces a roundish cluster shape that fits snugly between the walls. Hence the attainment of optimal cluster migration efficiency. The phenomenon of optimal confinement has also been confirmed for a larger cluster at *n* = 25.

**Figure 12:**
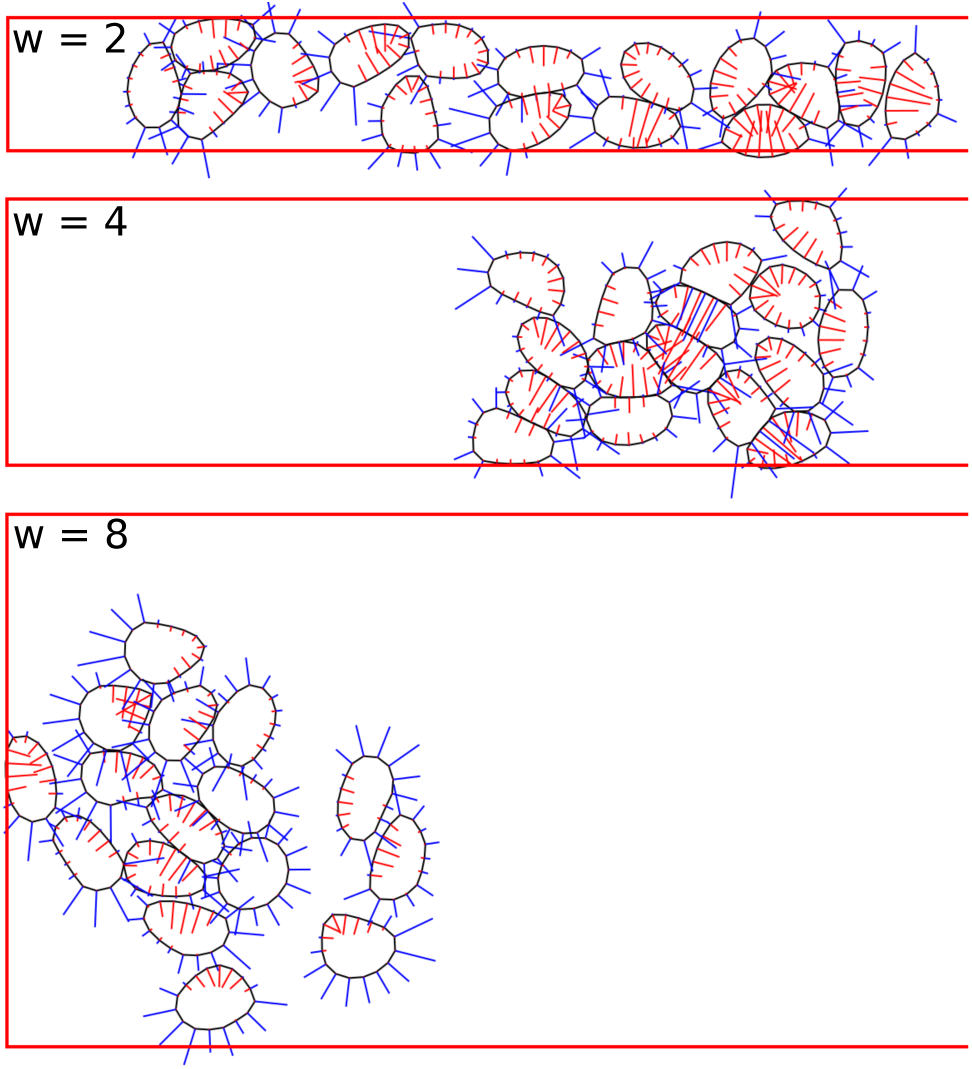
Snapshots of the migration of 16 cells in corridors of increasing width, *w* = 2, 4 and 8 at *t* = 225 min. Note the effect of over-confinement and under-confinement on the cluster configuration in the top and bottom panel, respectively. The mid-panel represents optimal confinement.

The concept of optimal corridor confinement was first proposed by Szabo *et al.* [14] based on a cellular Potts model. Besides, they collected *in vivo* data on NCC streams of different widths migrating in zebrafish and *Xenopus* embryos, and demonstrated that their cluster size and stream width conform to the idea of an optimal confinement. Our model confirms the existence of optimal confinement, and the optimal width identified here, 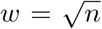, is consistent with both the *in vivo* and *in silico* data of Szabo *et al.* [14]. Furthermore, we have offered an explanation for the optimal confinement in terms of COA and CIL. Confinement tends to hinder COA’s ability to maintain coherence of the cluster, but it accentuates the role of CIL in keeping the cluster on a straight 1D path. An optimal confinement offers a balance between COA and CIL, both necessary ingredients for spontaneous collective migration.

We have also investigated the effect of initial cluster configuration by deviating from that of Fig. S5. For a fixed corridor width, say *w* = 4, we test initial configurations with the cells arranged into rectangular arrays of different width smaller than *w* positioned midway between the top and bottom walls, as well as initial configurations with the cells placed at random, non-overlapping positions (Fig. S7). The general trend is that the intensity of migration decreases for more elongated initial configurations, and that the random cluster has poorer migration intensity than the regular array (Fig. S8). As explained in the caption to Fig. S8, these observations are consistent with the understanding of confinement gained from Fig. 11.

## 4 Discussion

Our goal is to provide an explanation of spontaneous collective migration of neural crest cells in terms of key GTPases that regulate cell polarization and protrusion. From a broad perspective, we can summarize this work by the following three points.

First, we have adopted a modeling approach based on the biochemical signalling pathways as an alternative to the existing paradigms of continuum and agent-based modeling. Continuum models seek a “mean-field” description of the average orientation and movement of many cells without resolving the scales of individual cells [47–49]. However, it is challenging to represent intercellular interaction within the continuum formulation. Agent-based models address this concern by assigning rules on individual agents that recapitulate known cell-level behaviour [7, 13, 14, 50]. While helpful when little is known about the underlying biology, these models tend simply to give back the behaviour designed for. The biochemically-based model uses a finer level of resolution. Moreover, it seeks to connect cell-level behavior to intracellular signaling pathways that have been established experimentally. Thus, it can be more general and even simpler than models based on postulated rules. For example, our model predicts cell behaviours ranging from polarization to spontaneous collective migration from a few well-establish principles governing GTPases [22] and cell mechanics [51]. We should note that in recent years, biochemically-based modeling of cell mechanics has gained increasing interest and currency [52–54]. This is evidently motivated and enabled by growing experimental knowledge of the role of chemical signaling in complex behaviours such as collective migration [55–57].

Second, we have identified the persistence of polarity, or POP, as an essential factor in enabling spontaneous collective migration of cell clusters. Prior to this work, the extensive experimental studies of Mayor and coworkers had identified CIL and COA as key to spontaneous collective migration. Examining their recent models [13, 14], however, we have come to suspect that POP, an incidental feature of these models, may in fact be essential for the appearance of spontaneous collective migration. To test this hypothesis, we built a simpler model that recapitulates CIL and COA but with POP intentionally suppressed by erasing the cell polarity periodically and replacing it by a random “initial distribution” of the Rho GTPases. As illustrated in Fig. S2 and Movie 5, this model does not exhibit spontaneous collective migration in a corridor. Instead, the centroid of the cell cluster executes a 1D random walk once the cluster has moved away from the end of the corridor, where it enjoyed an initial directional migration thanks to the asymmetric initial condition.

Third, we have demonstrated that POP emerges naturally from the simultaneous action of COA and CIL. Thus, we have not only validated the existing proposal that COA and CIL cause spontaneous collective migration [6, 11], but also placed it on the concrete biological basis of Rac-Rho signaling. COA maintains the integrity of a cell cluster and ensures continual proximity and CIL interaction among neighbors. The two cooperate to suppress Rac1 and perpetuate an initial polarity and direction of migration that may have arisen from asymmetric boundary conditions and confinement. This is consistent with experimental observations that Rac1 suppression by drugs or Syndecan-4 enhances persistence in the polarity and motility in fibroblast and NCCs [31, 45, 46]. Thus, the model offers an explanation for the origin of POP; it is an emergent behavior based on COA and CIL rather than a separate mechanism to be postulated in addition to these two. This hypothesis can be tested by experiments that use drugs to partially suppress various proteins (or their activation levels) in the Rho GTPase signaling pathway [58, 59]. The resulting impairment of collective migration can be compared with model predictions of how COA, CIL and POP are affected by modulating the suitable Rac1 and RhoA rate constants.

It is interesting to put the current model in the context of directional motion and symmetry-breaking in vastly different systems, e.g. the active swimming of bacteria and flocking of insects or birds [17]. While alignment and directionality in macroscopic flocks rely on visual cues or pressure waves, we have shown that in the NCC context, it emerges from COA and CIL. Generally, one could view our POP as a device for alignment similar to the previously proposed directional persistence of random walk [7], persistence due to particle inertia [13], “persistence decay” for cell polarity [14] and collision-based alignment [60].

## 5 Conclusion

This paper presents a biochemistry-based model for the curious phenomenon of spontaneous collective migration of neural crest cells (NCCs). This represents an alternative to the paradigm of rule- or agent-based modeling. Starting from several well-established signaling pathways—autocatalytic activation and mutual inhibition of Rac1 and RhoA, Rac1 inhibition by contact and by membrane tension, and co-attraction from C3a/C3aR binding—as well as a standard vertex-based model for cell mechanics, we have predicted the following results:

- A single NCC polarizes with a protruding front featuring elevated Rac1 and a contracting rear featuring elevated RhoA. This, together with a randomized repolarization scheme, reproduces the tortuous trajectory of single-cell migration observed experimentally.
- In cell-cell encounters, the model reproduces the well-known contact inhibition of locomotion (CIL). The contact elevates the deactivation rate of Rac1 and the activation rate of RhoA, thereby neutralizing the protrusion front and turning it into a retracting rear.
- The model recapitulate the effect of co-attraction (COA) in preventing a cluster of cells from scattering due to CIL. COA thus provides an aggregative effect that balances the dispersal effect of CIL to keep the integrity of cell clusters.
- In a confining corridor, the simultaneous action of COA and CIL produces a persistence of cell polarity (POP), such that cell clusters will continue to migrate over great lengths in the direction favored by the initial configuration. The collective migration is more robust for larger clusters. For the model parameters used, the predicted cluster migration speed and persistence agree with experiments.
- There exists an optimal confinement that produces maximum persistence and speed of collective migration. For a cluster of *n* cells, the optimal corridor width is 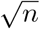 times the single cell diameter. This is consistent with prior *in vivo* and *in silico* data.

These results confirm the current hypothesis that the spontaneous collective migration of NCCs is due to CIL and COA, and provide a more complete picture by elucidating the biological basis for POP.

We must emphasize the limitations of this work as well. Collective migration of NCCs is a highly complex process, with multiple potential mechanisms at work for cranial and trunk NCCs and in different species [5, 61]. This study is restricted to a relatively simple *in vitro* scenario of NCCs spontaneously migrating down a corridor in the absence of chemoattractants. Our model does not preassign “leader” or “trailblazer” cells [2], and as such is likely to be relevant only to cranial NCCs that appear to possess a homogeneous migratory capability [5]. Keeping in mind the complexity of *in vivo* NCC migration, one may adapt the present theoretical framework to explore certain aspects of that phenomenon. A first step in this direction will be to add a chemokine such as Sdf1 and study how it modifies the CIL-COA-POP mechanism during chemotaxis [3, 38].

On a more technical level, the model incorporates numerous simplifications. For example, cellcell adhesion is not explicitly accounted for, nor is the actomyosin apparatus for force generation. Arguably, the role of N-cadherins has been subsumed into the CIL pathway [38]. But cell-cell adhesion may also directly supplement COA in maintaining the integrity of the cluster. Similarly, cell-substrate or cell-ECM adhesion is neglected. In reality, protrusion and migration are enabled by the actomyosin contractile apparatus that is anchored onto the substrate or ECM. The model treats this in a phenomenological way via the protrusion/contraction force parameters (Eq. 11) and the frictional factor (Eq. 9). Finally, COA is modeled by summing up C3a signals from individual cells in a linearly additive manner. In reality, response to C3a-C3aR binding probably saturates nonlinearly. For lack of experimental data, the model has adopted the simplest reasonable form. The validity of these simplifications remains to be verified by carefully designed experiments that isolate the factors in question.

## Acknowledgement

The authors acknowledge financial support by the Natural Sciences and Engineering Research Council of Canada. We also thank Philip Maini and Roberto Mayor for discussions and comments on various aspects of the project.

## References

[1] R. McLennan, L. Dyson, K. W. Prather, J. A. Morrison, R. E. Baker, P. K. Maini, P. M. Kulesa, Multiscale mechanisms of cell migration during development: theory and experiment, Development 139 (16) (2012) 2935–2944.

[2] R. McLennan, L. J. Schumacher, J. A. Morrison, J. M. Teddy, D. A. Ridenour, A. C. Box, C. L. Semerad, H. Li, W. McDowell, D. Kay, P. K. Maini, R. E. Baker, P. M. Kulesa, Vegf signals induce trailblazer cell identity that drives neural crest migration, Developmental biology 407 (1) (2015) 12–25.

[3] A. Shellard, R. Mayor, Chemotaxis during neural crest migration, Semin. Cell Dev. Biol. 55 (2016) 111–118.

[4] A. Szabó, R. Mayor, Modelling collective cell migration of neural crest, Curr. Opin. Cell Biol. 42 (2016) 22–28.

[5] J. Richardson, A. Gauert, L. B. Montecinos, L. Fanlo, Z. M. Alhashem, R. Assar, E. Marti, A. Kabla, S. Härtel, C. Linker, Leader cells define directionality of trunk, but not cranial, neural crest cell migration, Cell Rep. 15 (9) (2016) 2076–2088.

[6] C. Carmona-Fontaine, E. Theveneau, A. Tzekou, M. Tada, M. Woods, K. M. Page, M. Parsons, J. D. Lambris, R. Mayor, Complement fragment c3a controls mutual cell attraction during collective cell migration, Dev. Cell 21 (6) (2011) 1026–1037.

[7] S. Huang, C. Brangwynne, K. Parker, D. Ingber, Symmetry-breaking in mammalian cell cohort migration during tissue pattern formation: Role of random-walk persistence, Cytoskeleton 61 (4) (2005) 201–213.

[8] S. R. K. Vedula, M. C. Leong, T. L. Lai, P. Hersen, A. J. Kabla, C. T. Lim, B. Ladoux, Emerging modes of collective cell migration induced by geometrical constraints, Proc. Natl. Acad. Sci. U.S.A. 109 (32) (2012) 12974–12979.

[9] E. F. Boer, E. D. Howell, T. F. Schilling, C. A. Jette, R. A. Stewart, Fascin1-dependent filopodia are required for directional migration of a subset of neural crest cells, PLoS Genet. 11 (1) (2015) e1004946.

[10] A. J. Burns, J.-M. M. Delalande, N. M. Le Douarin, In ovo transplantation of enteric nervous system precursors from vagal to sacral neural crest results in extensive hindgut colonisation, Development 129 (12) (2002) 2785–2796.

[11] C. Carmona-Fontaine, H. K. Matthews, S. Kuriyama, M. Moreno, G. A. Dunn, M. Parsons, C. D. Stern, R. Mayor, Contact inhibition of locomotion in vivo controls neural crest directional migration, Nature 456 (7224) (2008) 957–961.

[12] R. Mayor, E. Theveneau, The role of the non-canonical wnt–planar cell polarity pathway in neural crest migration, Biochem. J 457 (1) (2014) 19–26.

[13] M. L. Woods, C. Carmona-Fontaine, C. P. Barnes, I. D. Couzin, R. Mayor, K. M. Page, Directional collective cell migration emerges as a property of cell interactions, PloS One 9 (9) (2014) e104969.

[14] A. Szabó, M. Melchionda, G. Nastasi, M. L. Woods, S. Campo, R. Perris, R. Mayor, In vivo confinement promotes collective migration of neural crest cells, J. Cell Biol. 213 (5) (2016) 543–555.

[15] A. Szabó, R. Ünnep, E. Méhes, W. O. Twal, W. S. Argraves, Y. Cao, A. Czirók, Collective cell motion in endothelial monolayers, Phys. Biol. 7 (2010) 046007.

[16] T. Vicsek, A. Czirók, E. Ben-Jacob, I. Cohen, O. Shochet, Novel type of phase transition in a system of self-driven particles, Phys. Rev. Lett. 75 (1995) 1226–1229.

[17] M. C. Marchetti, J. F. Joanny, S. Ramaswamy, T. B. Liverpool, J. Prost, M. Rao, R. A. Simha, Hydrodynamics of soft active matter, Rev. Mod. Phys. 85 (2013) 1143–1189.

[18] A. Roycroft, R. Mayor, Molecular basis of contact inhibition of locomotion, Cell. Mol. Life Sci. 73 (6) (2016) 1119–1130.

[19] B. Vanderlei, J. J. Feng, L. Edelstein-Keshet, A computational model of cell polarization and motility coupling mechanics and biochemistry, Multiscale Model. Simul. 9 (4) (2011) 1420–1443.

[20] M. P. Neilson, J. A. Mackenzie, S. D. Webb, R. H. Insall, Modelling cell movement and chemotaxis using pseudopod-based feedback, SIAM J. Sci. Comput. 33 (3) (2011) 1035–1057.

[21] M. Raftopoulou, A. Hall, Cell migration: Rho GTPases lead the way, Dev. Biol. 265 (1) (2004) 23–32.

[22] A. B. Jaffe, A. Hall, Rho GTPases: biochemistry and biology, Annu. Rev. Cell Dev. Biol. 21 (2005) 247–269.

[23] M. M. Zegers, P. Friedl, Rho GTPases in collective cell migration, Small GTPases 5 (3) (2014) e983869.

[24] A. J. Ridley, Rho GTPase signalling in cell migration, Curr. Opin. Cell Biol. 36 (2015) 103–112.

[25] R. Mayor, S. Etienne-Manneville, The front and rear of collective cell migration, Nat. Rev. Mol. Cell Biol. 17 (2) (2016) 97–109.

[26] D. V. Köster, S. Mayor, Cortical actin and the plasma membrane: inextricably intertwined, Curr. Opin. Cell Biol. 38 (2016) 81–89.

[27] A. G. Fletcher, M. Osterfield, R. E. Baker, S. Y. Shvartsman, Vertex models of epithelial morphogenesis, Biophys. J. 106 (2014) 2291–2304.

[28] H. Lan, Q. Wang, R. Fernandez-Gonzalez, J. J. Feng, A biomechanical model for cell polarization and intercalation during drosophila germband extension, Phys. Biol. 12 (2015) 056011.

[29] Y. Mori, A. Jilkine, L. Edelstein-Keshet, Wave-pinning and cell polarity from a bistable reaction-diffusion system, Biophys. J. 94 (9) (2008) 3684–3697.

[30] B. Huang, M. Lu, M. K. Jolly, I. Tsarfaty, J. Onuchic, E. Ben-Jacob, The three-way switch operation of Rac1/RhoA GTPase-based circuit controlling amoeboid-hybrid-mesenchymal transition, Sci. Rep. 4 (2014) 6449.

[31] R. Pankov, Y. Endo, S. Even-Ram, M. Araki, K. Clark, E. Cukierman, K. Matsumoto, K. M. Yamada, A Rac switch regulates random versus directionally persistent cell migration, J. Cell Biol. 170 (5) (2005) 793–802.

[32] A. R. Houk, A. Jilkine, C. O. Mejean, R. Boltyanskiy, E. R. Dufresne, S. B. Angenent, S. J. Altschuler, L. F. Wu, O. D. Weiner, Membrane tension maintains cell polarity by confining signals to the leading edge during neutrophil migration, Cell 148 (1) (2012) 175–188.

[33] A. Diz-Muñoz, K. Thurley, S. Chintamen, S. J. Altschuler, L. F. Wu, D. A. Fletcher, O. D. Weiner, Membrane tension acts through pld2 and mtorc2 to limit actin network assembly during neutrophil migration, PLoS Biol. 14 (6) (2016) e1002474.

[34] W. R. Holmes, L. Edelstein-Keshet, Analysis of a minimal Rho-GTPase circuit regulating cell shape, Phys. Biol. 13 (4) (2016) 046001.

[35] R. J. Petrie, A. D. Doyle, K. M. Yamada, Random versus directionally persistent cell migration, Nat. Rev. Mol. Cell Biol. 10 (8) (2009) 538–549.

[36] M. Krause, A. Gautreau, Steering cell migration: lamellipodium dynamics and the regulation of directional persistence, Nat. Rev. Mol. Cell Biol. 15 (9) (2014) 577–590.

[37] A. A. Potdar, J. Lu, J. Jeon, A. M. Weaver, P. T. Cummings, Bimodal analysis of mammary epithelial cell migration in two dimensions, Ann. Biomed. Eng. 37 (1) (2009) 230–245.

[38] E. Theveneau, L. Marchant, S. Kuriyama, M. Gull, B. Moepps, M. Parsons, R. Mayor, Collective chemotaxis requires contact-dependent cell polarity, Dev. Cell 19 (1) (2010) 39–53.

[39] R. Gorelik, A. Gautreau, Quantitative and unbiased analysis of directional persistence in cell migration, Nat. Protoc. 9 (8) (2014) 1931–1943.

[40] R. Moore, E. Theveneau, S. Pozzi, P. Alexandre, J. Richardson, A. Merks, M. Parsons, J. Kashef, C. Linker, R. Mayor, Par3 controls neural crest migration by promoting microtubule catastrophe during contact inhibition of locomotion, Development 140 (23) (2013) 4763–4775.

[41] E. Scarpa, A. Roycroft, E. Theveneau, E. Terriac, M. Piel, R. Mayor, A novel method to study contact inhibition of locomotion using micropatterned substrates, Biol. Open 2 (9) (2013) 901–906.

[42] S. Vedel, S. Tay, D. M. Johnston, H. Bruus, S. R. Quake, Migration of cells in a social context, Proc. Natl. Acad. Sci. U.S.A. 110 (1) (2013) 129–134.

[43] E. A. Novikova, M. Raab, D. E. Discher, C. Storm, Persistence-driven durotaxis: Generic, directed motility in rigidity gradients, Phys. Rev. Lett. 118 (7) (2017) 078103.

[44] M. F. Ware, A. Wells, D. A. Lauffenburger, Epidermal growth factor alters fibroblast migration speed and directional persistence reciprocally and in a matrix-dependent manner, J. Cell Sci. 111 (16) (1998) 2423–2432.

[45] M. D. Bass, K. A. Roach, M. R. Morgan, Z. Mostafavi-Pour, T. Schoen, T. Muramatsu, U. Mayer, C. Ballestrem, J. P. Spatz, M. J. Humphries, Syndecan-4–dependent Rac1 regulation determines directional migration in response to the extracellular matrix, J. Cell Biol. 177 (3) (2007) 527–538.

[46] H. K. Matthews, L. Marchant, C. Carmona-Fontaine, S. Kuriyama, J. Larraín, M. R. Holt, M. Parsons, R. Mayor, Directional migration of neural crest cells in vivo is regulated by Syndecan-4/Rac1 and non-canonical Wnt signaling/RhoA, Development 135 (10) (2008) 1771–1780.

[47] J. Löber, F. Ziebert, I. S. Aranson, Collisions of deformable cells lead to collective migration, Sci. Rep. 5 (2015) 9172.

[48] S. Najem, M. Grant, Phase-field model for collective cell migration, Phys. Rev. E 93 (2016) 052405.

[49] W. Marth, S. Praetorius, A. Voigt, A mechanism for cell motility by active polar gels, J. R. Soc. Interface 12 (107) (2015) 20150161.

[50] B. Smeets, R. Alert, J. Pešek, I. Pagonabarraga, H. Ramon, R. Vincent, Emergent structures and dynamics of cell colonies by contact inhibition of locomotion, Proc. Natl. Acad. Sci. U.S.A. 113 (51) (2016) 14621–14626.

[51] T. Wu, J. J. Feng, Modeling the mechanosensitivity of neutrophils passing through a narrow channel, Biophys. J. 109 (11) (2015) 2235–2245.

[52] B. A. Camley, J. Zimmermann, H. Levine, W.-J. Rappel, Collective signal processing in cluster chemotaxis: Roles of adaptation, amplification, and co-attraction in collective guidance, PLoS Comput. Biol. 12 (7) (2016) e1005008.

[53] J. Varennes, B. Han, A. Mugler, Collective chemotaxis through noisy multicellular gradient sensing, Biophys. J. 111 (3) (2016) 640–649.

[54] J. Delile, M. Herrmann, N. Peyriéras, R. Doursat, A cell-based computational model of early embryogenesis coupling mechanical behaviour and gene regulation, Nat. Commun. 8 (2017) 13929.

[55] F. Fagotto, The cellular basis of tissue separation, Development 141 (17) (2014) 3303–3318.

[56] C. Collins, W. J. Nelson, Running with neighbors: coordinating cell migration and cell–cell adhesion, Curr. Opin. Cell Biol. 36 (2015) 62–70.

[57] T. Das, K. Safferling, S. Rausch, N. Grabe, H. Boehm, J. P. Spatz, A molecular mechanotransduction pathway regulates collective migration of epithelial cells, Nat. Cell Biol. 17 (3) (2015) 276–287.

[58] J. Park, W. R. Holmes, S. H. Lee, H.-N. Kim, D.-H. Kim, M. K. Kwak, C. J. Wang, L. Edelstein-Keshet, A. Levchenko, Mechanochemical feedback underlies coexistence of qualitatively distinct cell polarity patterns within diverse cell populations, Proc. Natl. Acad. Sci. U.S.A. 114 (28) (2017) E5750–E5759.

[59] W. R. Holmes, J. Park, A. Levchenko, L. Edelstein-Keshet, A mathematical model coupling polarity signaling to cell adhesion explains diverse cell migration patterns, PLoS Comput. Biol. 13 (5) (2017) e1005524.

[60] L. Coburn, L. Cerone, C. Torney, I. D. Couzin, Z. Neufeld, Tactile interactions lead to coherent motion and enhanced chemotaxis of migrating cells, Phys. Biol. 10 (4) (2013) 046002.

[61] P. M. Kulesa, R. McLennan, Neural crest migration: trailblazing ahead, F1000Prime Rep. 7 (2015) 02.

